# UDP-glucose:anthocyanidin 3-*O*-glucoside-2”-*O*-glucosyltransferase catalyzes further glycosylation of anthocyanins in purple *Ipomoea batatas*

**DOI:** 10.1101/332486

**Authors:** Hongxia Wang, Chengyuan Wang, Weijuan Fan, Jun Yang, Ingo Appelhagen, Yinliang Wu, Peng Zhang

## Abstract

Glycosylation contributes to the diversity and stability of anthocyanins in plants. The process is catalyzed by various glucosyltransferases using different anthocyanidin aglycones and glycosyl donors. An anthocyanidin 3*-O-*glucoside-2”*-O-*glucosyltransferase (3GGT) from purple sweetpotato (cv. Ayamurasaki) served for the catalytic conversion of anthocyanidin 3*-O-*glucoside into anthocyanidin 3*-O-*sophoroside, which is functionally different from the 3GGT ortholog of *Arabidopsis*. The phylogenetic analysis indicates regioselectivity of 3GGT using UDP-xylose or UDP-glucose as the glycosyl is divergent between *Convolvulaceae* and *Arabidopsis*. Homology-based protein modeling and site-directed mutagenesis of Ib3GGT and At3GGT suggested that the Thr-138 of Ib3GGT is a key amino acid residue for UDP-glucose recognition and plays a major role in sugar donor selectivity. The wild type and *ugt79b1* mutants of *Arabidopsis* plants overexpressing *Ib3GGT* produced the new component cyanidin 3*-O-*sophoroside. Moreover, *Ib3GGT* expression was associated with anthocyanin accumulation in different tissues during Ayamurasaki plant development and was regulated by the transcription factor IbMYB1. The localization assay of Ib3GGT showed that further glycosylation occurs in the cytosol and not endoplasmic reticulum. The present study revealed the function of Ib3GGT in further glycosylation of anthocyanins and its Thr-138 is the key amino acid residue for UDP-glucose recognition.

## Introduction

Anthocyanins are major secondary metabolites responsible for color variation in plants, exhibiting health-promoting properties (de Pascual-Teresa and Sanchez-Ballesta, 2008; He and Giusti, 2010). The basic structures of anthocyanins are mono- and di-glycosylated forms in common anthocyanidins, which include cyanidin, delphinidin, malvidin, pelargonidin, peonidin, and petunidin (Moglia et al., 2014). Different sugar moieties, i.e., glucose, galactose, xylose, arabinose, or fructose can be linked to hydroxyl groups at 3, 5, 7, 3’, and 5’ positions, with the glycosylation at the 3^rd^ position on the C-ring ubiquitously (Andersen and Jordheim, 2010). Glycosylation of 3-OH is catalyzed by a series of UDP carbohydrate-dependent glycosyltransferases (UGTs), which utilize the nucleotide-activated sugars as donor substrates and anthocyanidin aglycones or anthocyanins as acceptors. These activities increase the structural diversity of anthocyanins by adding different types and/or numbers of sugar moieties on various positions (Gachon et al., 2005). The glycosylation of anthocyanin is speculated to occur on the cytoplasmic surface of the endoplasmic reticulum (ER), and may serve as a signal for the transport of anthocyanins to vacuoles via multiple pathways; this transport is essential for the stable storage of anthocyanins in vacuoles (Ono et al., 2006; Matsuba et al., 2010; Sun et al., 2012; Zhao et al., 2011; Zhao, 2015). Glycosylation also participates in the fine adjustment and stabilization of flower pigmentation in ornamental plants (Yonekura-Sakakibara et al., 2012).

Monoglycosylation of anthocyanidins produces anthocyanidin 3-*O*-glucosides, the first major stable colored pigments in the anthocyanin biosynthesis pathway (Griesser et al., 2008a; Montefiori et al., 2011). Deficiency of the activity of the corresponding UDP-glucose:flavonoid 3*-O-*glycosyltransferase (UF3GT), in maize *bronze1* and *Arabidopsis anl1*, results in a significantly suppressed accumulation of anthocyanin (Fedoroff et al., 1984; Kubo et al., 2007). Until now, UF3GT is one of the well-characterized UGTs related to anthocyanin biosynthesis (Gachon et al., 2005; Yonekura-Sakakibara and Hanada, 2011). Further glycosylation of anthocyanidin 3*-O-*glucosides involves diverse sugars in different species, such as UDP-rhamnose, UDP-glucose, UDP-xylose, and UDP-arabinose, as donor substrates to be added at a species-specific position to the glycosides of mono 3-*O*-glycosylated anthocyanins (Yonekura-Sakakibara et al., 2012). The mutants affected in this further glycosylation function may be impacted in the anthocyanin accumulation, as reported in petunia and Japanese morning glory (Kroon et al., 1994; Morita et al., 2005). Since all the UGT proteins are highly similar in their secondary and tertiary structures, with a defined fold structure and highly conserved putative secondary product glycosyltransferase (PSPG) motif (Breton et al., 2006; Lairson et al., 2008; Osmani et al., 2009), structure-based modeling have identified the key residues of UF3GT responsible for sugar donor specificity (Kubo et al., 2004) in *Arabidopsis* (Kim et al., 2013), *Freesia hybrid* (Sun et al., 2016), grapes (Offen et al., 2006; Ono et al., 2010), lamiales (Noguchi et al., 2009), perilla (Noguchi et al., 2009), and red daisy (Osmani et al., 2009). Nevertheless, the residues involved in UDP-sugar selectivity in 3GGT are yet unknown.

Although the anthocyanidin decoration by glycosylation is progressive, it commonly begins with 3-*O*-glycosylation to ensure the stability of the aglycon. Additional glycosylation leads to the compound and functional diversity, thereby contributing to several varieties of anthocyanins in the plant (Gachon et al., 2005; Caputi et al., 2012). To date, more than 600 anthocyanins or their derivatives have been identified in nature (Glover and Martin, 2012); however, only a limited number of genes encoding UFGTs in different species have been well characterized. Several flavonoid 3*-O-*glycosyltransferases have been characterized in *Arabidopsis* (Kubo et al., 2007; Saito et al., 2013), strawberry (Griesser et al., 2008a, 2008b), grapes (Offen et al., 2006), and maize (Fedoroff et al., 1984). In addition, for further flavonoid glycosylation multiple UGTs were also characterized, including anthocyanidin 3*-O-*glucoside 6”*-O-*rhamnosyltransferase in *Petunia hybrida* (Kroon et al., 1994), anthocyanidin 3*-O-*glucoside 2”*-O-*glucuronosyltransferase in red daisy flowers (Sawada et al., 2005), and flavonol 3*-O-*glucoside 2”*-O-*glucosyltransferase in *Arabidopsis* (Yonekura-Sakakibara et al., 2014). Apparently, the divergence towards different glycosylation types occurs at this step. At the same 2” position, different glycosylation types, i.e., glycosylation or xylosylation, are found in various plant species. In morning glory, anthocyanidin 3*-O-*glucoside 2”*-O-*glucosyltransferase catalyzes the addition of a glucose molecule to anthocyanidin 3*-O-*glucosides on the 2” position to form anthocyanidin 3*-O-*sophorosides (Morita et al., 2005). In *Arabidopsis*, further glycosylation of the 3-*O*-glucoside is catalyzed by anthocyanidin 3*-O-*glucoside 2”*-O-*xylosyltransferase (AtA3G2XylT, i.e., At3GGT) to add one xylose molecule specifically to the first glucose residue (Yonekura-Sakakibara et al., 2012). Further decorations, for example, to the diversity or functionality, malonylation and aromatic acylation rely on the glycosylation of anthocyanidins (Sasaki et al., 2014).

Purple sweetpotato (*Ipomoea batatas*) accumulates a lot of anthocyanins in storage roots. Anthocyanidin 3*-O-*glucoside-2”*-O-*glucoside (anthocyanin 3*-O-*sophoroside) and derivatives are the major anthocyanin compounds (Tian et al., 2005). So far, at least 26 components, mostly caffeoylated, coumarylated or feruloylated anthocyanidin glucosides have been identified (Truong et al., 2009; Lee et al., 2013). In contrast, 11 anthocyanins have been identified in *Arabidopsis*; all of them derived from cyanidin 3*-O-*glucoside-2”*-O-*xyloside (Tohge et al., 2005; Yonekura-Sakakibara et al., 2012; Kovinich et al., 2014). Therefore, unlike *Arabidopsis*, purple sweetpotato uses UDP-glucose as sugar donor for further glycosylation of anthocyanidin 3*-O-*glucosides to form anthocyanidin 3*-O-*sophorosides. In the present study, we characterized a UFGT, termed as UDP-glucose:anthocyanidin 3*-O-*glucoside-2”*-O-*glucosyltransferase (IbA3G2GluT, i.e., Ib3GGT) that catalyzes the anthocyanin glycosylation in purple sweetpotato and its key amino acid for sugar donor selectivity.

## Materials and methods

### Plant materials

The purple-fleshed sweetpotato (*Ipomoea batatas* Lam.) cultivar Ayamurasaki was used in this study. The *in vitro* shoot cultures were subcultured on SBM medium (MS salts including vitamins + 0.3 mg/L VB1 + 30 g/L sucrose, pH 5.8) in plant growth chambers under a 16 h photoperiod provided by cool-white fluorescent tubes (∼50 μmol/m^2^/s), at 25°C and 50% relative humidity. One-month-old plantlets were transplanted into plastic pots containing well-mixed soil (soil:peat:perlite, 1:1:1) and grown in the greenhouse (16 h/8 h light/dark cycle, 25°C day/night). Various tissues including leaves, stems, fibrous roots, and storage roots of sweetpotato plants at different developmental stages were harvested from the pot- or field-grown plants for multiple analyses. The *Arabidopsis* plants were grown under a 16 h/8 h light/dark cycle, at 22°C in the growth chamber.

### Plasmid construction and production of transgenic sweetpotato

The open reading frame of *Ib3GGT* (1380 bp) was amplified from the cDNA of sweetpotato Ayamurasaki using the primers Ib3GGTF (5’-CGGGGTACCATGGGTTCTCAAGCAACAAC-3’, KpnI site underlined) and Ib3GGTR (5’-AATGTCGACTCATCCAAGGAGATCCTGCA-3’, SalI site underlined). This fragment was inserted into the KpnI/SalI sites of the pCAMBIA1301-based plant expression vector to generate the binary vector pOE-Ib3GGT containing the expression cassette of *Ib3GGT* driven by the CaMV 35S promoter. The pRNAi-Ib3GGT binary vector was manipulated to express double-stranded hairpin RNA of the 252 bp *Ib3GGT* fragment (382–633 bp) based on the pRNAi-DFR vector (Wang et al., 2013). Then, pOE-Ib3GGT and pRNAi-Ib3GGT were introduced into *Agrobacterium tumefaciens* strain LBA4404 for sweetpotato transformation, as described previously (Yang et al., 2011). Transgenic plants were produced and verified for *Ib3GGT* expression by real-time RT-PCR. For total *Ib3GGT* expression, an internal primer pair of *Ib3GGT* was designed for detecting the *Ib3GGT* expression in WT, OE-Ib3GGT and RNAi-Ib3GGT plants by real-time RT-PCR (Table S2). The *Actin* gene of sweetpotato was used as an internal control for gene amplification.

### Transformation and analysis of *Ib3GGT*-overexpressing *Arabidopsis*

Two independent UGT79B1 *Arabidopsis* transposon mutants, *ugt79b1-1* and *ugt79b1-2* (Kuromori et al., 2004; Ito et al., 2005), along with the WT Nossen and ecotype Col-0 were transformed with *A. tumefaciens* LB4404 harboring pOE-Ib3GGT, using the floral dip method (Clough and Bent, 1998). The transformants were selected on 1/2 MS medium containing 50 mg/L hygromycin for Nossen and mutants or 25 mg/L hygromycin for Col-0 plants. The RNA extracted from T3 homozygous *Arabidopsis* seedlings was used for RT-PCR analysis. The primer pairs used to detect the expression of *At3GGT* and *Ib3GGT* in WT and transgenic *Arabidopsis* plants were designed using software Primer 3.0 and listed in Table S2. *At3GGT* were amplified a 223-bp fragment from position +369 to +591 bp and *Ib3GGT* were amplified a 189-bp fragment from +1009 to +1197 bp. The *Actin* gene of *Arabidopsis* was used as a reference gene.

### Phylogenetic Analysis

To construct a phylogenetic tree, 16 UGT protein sequences obtained from NCBI GenBank were aligned by ClustalW and implemented in MEGA6 (Tamura et al., 2013). Ten closely related UGTs were used to illustrate the relationship. The maximum likelihood method was used to obtain the alignment results (Stamatakis, 2014). Bootstrap values were obtained with 1000 replications.

### Site-directed mutagenesis and *in vitro* enzymatic assay of recombinant Ib3GGT and At3GGT

The full-length sequence of the *Ib3GGT* gene was amplified by PCR using the primers IbGGT-FP (5’-CCC<u>AAGCTT</u>ATGGGTTCTCAAGCAACAAC-3’, HindIII site underlined) and IbGGT-RP (5’-CGC*GGATCC*TCA<u>CATCACCATCACCATCAC</u>TCCAAGGAGATCCTGCA-3’, BamHI site and 6 His sites underlined). The full-length *At3GGT* was amplified by PCR using the primers AtGGT-FP (5’-GG<u>GGTACCA</u>TGGGTGTTTTTGGATCGAA-3’, KpnI site underlined) and AtGGT-RP (5’-CG<u>GAATTC</u>TCA<u>CATCACCATCACCATCAC</u>TGACTTCACAAGTTCAATTA AATT-3’, EcoRI site and 6 His sites underlined). Site-directed mutations were generated by changing the Thr-138 nucleotide ACC into ATT in *Ib3GGT* and Ile-142 ATC into ACT in *At3GGT* using PCR-based amplification with a Phusion Site-Directed Mutagenesis Kit (Thermo Scientific). The sequence fragments, with or without the mutation of the 3GGTs, were cloned into the pYES2 vector and introduced in *Saccharomyces cerevisiae* BY4742 according to the manufacturer’s instructions (Cat# V825-20, Invitrogen). The recombinant 3GGT proteins were induced by replacing the carbon source from 2% glucose to 2% galactose in the SC-U medium. The reaction mixture for the 3GGT enzymatic assay consisted of 100 mM phosphate buffer (pH 7.0), 0.6 mM flavonoid aglycones (cyanidin, cyanidin3*-O-*glucoside, cyanidin 3,5*-O-*diglucoside, or flavonol 3*-O-*glucoside), 1 mM UDP-glucose, and 20 μL of crude yeast extract as the enzymatic solution in a reaction volume of 100 μL. After incubation for 2 h at 37°C, the reaction was terminated by centrifugation. The enzymatic activity of mutant 3GGT was assessed by cyanidin 3-*O*-glucoside as the acceptor substrate and different UDP-sugars (UDP-glucose, UDP-xylose, UDP-galactose or UDP-arabinose) as a sugar donor.

### LC-MS analyses of metabolites obtained by enzymatic reaction

Ten µL of filtered supernatants were analyzed on an Agilent HPLC1200-MSD/Q-TOF 6520 system (Agilent, Waldbronn, Germany) as described previously (Wang et al., 2013). Briefly, the mobile phase consisted of 0.5% (v/v) acetic acid in water (eluent A) and 100% acetonitrile (eluent B). The samples eluted at a flow rate of 0.2 mL/min passed through a reverse-phase C18 column (Agilent ZORBAX Eclipse XDB, 4.6 × 50 mm ID, 1.8 μm), and a DAD detector at 530 nm monitored the anthocyanin. Subsequently, an ESI interfaced Q-TOF mass detector (*m/z* 40–1500) collected the mass m/z data that were processed by Agilent Mass Hunter Qualitative Analysis (version 3.0) for the estimation of accurate molecular mass as well as spectrum evaluation. Cyanidin 3-*O*-sophoroside (Tongtian, Shanghai, China) was used as a standard.

### Subcellular localization of Ib3GGT in plant cells

The *Ib3GGT* gene was amplified by PCR using Pfu polymerase (Takara, Shanghai, China) to obtain a non-stop coding sequence using the primers FPGGT_L (5’-AATGTCGACATGGGTTCTCAAGCAACAAC-3’, SalI site underlined) and RPGGT_L (5’-GGACTAGTCCAAGGAGATCCTGCAGTT-3’, SpeI site underlined).

Ib3GGT-eGFP was constructed by inserting the *Ib3GGT* fragment into the corresponding sites of a modified pCambia1300 to fuse with the eGFP coding sequence. The construction of the ER-marker (Nelson et al., 2007) and the expression construct for mRFP (Claudia et al., 2017) has been described elsewhere. The ER marker, ER-mCherry, contains a signal peptide of AtWAK2 at the N-terminal and a synthetic HDEL at the C-terminal (He et al., 1999; Nelson et al., 2007). All constructs were introduced into *A. tumefaciens* GV3101 (pMP90). The growth conditions for *N. benthamiana* and *A. tumefaciens*, as well as the agro-infiltration procedure, were described previously (Leuzinger et al., 2013). The images were acquired 36 h post-infiltration with a Leica SP8X confocal microscope equipped with a Leica HC PL APO CS2 63x/1.20 water immersion objective. The GFP fluorescence was detected by hybrid detector HyD1 in the range of 500–540 nm and excited using the 488-nm line of an argon ion laser. mCherry and mRFP fluorescence were detected in the range of 580–630 nm by HyD2 after excitation at 561 nm with a diode-pumped solid-state laser. Both fluorophores were recorded line-by-line sequentially at a 3-to 4-fold average in a background noise-dependent manner. The Leica Application Suite X software was used for image acquisition and intensity estimations.

### Anthocyanin measurement and detection

Total anthocyanins in the WT and transgenic lines were extracted using previously described methods with slight modifications (Wang et al., 2013). The total content of anthocyanin in the WT and transgenic lines was quantified as cyanidin 3-*O*-sophoroside equivalent. The anthocyanin autofluorescence in epidermal cells of sweetpotato leaves was examined using a PCM-2000/Nikon Eclipse 600 laser-scanning microscope (Nikon, Japan) equipped with an argon and helium-neon laser (excitation 488 nm, emission 544 nm).

### Luciferase assay

The *Ib3GGT* promoter (2000 bp) was amplified by the primers Ib3GGTprFP (3’-AACTGCAGTTCAGTCAGGCAATCACAGG-5’, PstI site underlined) and Ib3GGTprRP (3’-CGCGGATCCAATAATACCTAGCTAGCT-5’, BamHI site underlined) and cloned into the pLL00R vector to generate the luciferase reporter vector. The *IbMYB1* gene was amplified by the primers IbMYB1FP (3’-GGGGTACCATGGTTATTTCATCTGTATG, KpnI site underlined) and IbMYB1RP (3’-AACTGCAGTTAGCTTAACAGTTCTGAC-5’, PstI site underlined) and subcloned into pCAMBIA1300 to generate the CaMV35S-IbMYB1 effector plasmid. *A. tumefaciens* strain GV3101 harboring the *Ib3GGT* promoter-LUC reporter and CaMV 35S-IbMYB1 effector was infiltrated into the 5-week-old *N. benthamiana* leaves using a needleless syringe for assessing the luciferase activity. The plants were grown for 48 h (16 h/8 h light/dark cycle, 25°C day/night), followed by injecting the leaves with 0.94 mM luciferin as substrate. The leaves were collected in the dark after 3 min and luciferase signals detected on a Tanon-5200 image system. The LUC reporter empty vector with 35S-IbMYB1 or *Ib3GGT* promoter-LUC reporter with empty effector vector was also co-infiltrated as a negative control. These experiments were repeated at least three times, and similar results were obtained.

### Molecular modeling of Ib3GGT and At3GGT active sites

The 3D models of Ib3GGT and At3GGT were generated using SWISS-MODEL workspace (Biasini et al., 2014; Wetterhorn et al., 2016) and I-TASSER server (Hiromoto et al., 2006) based on the structure of *N*-/*O*-glucosyltransferase of *A. thaliana* that served as a template (UGT72B1 PDB ID: 2VCE, Brazier-Hicks et al., 2007). The substrate binding sites were predicted by superposing both models to UGT72B1 using the COOT program (Emsley et al., 2010).

### Statistical analyses

All data were represented as mean ± SD from at least three biological replicates. One-way ANOVA analyses were performed by using SPSS Statistics 17.0 to Duncan’s multiple comparison tests. A value of *P* < 0.05 was considered as statistically significant difference.

## Results

### Comparison of anthocyanins indicates different further glycosylation patterns in sweetpotato and *Arabidopsis*

In purple sweetpotato cv. Ayamurasaki, anthocyanins include aromatically acylated anthocyanidin 3*-O-*sophoroside and derivatives, whereas in *Arabidopsis* Col-0, anthocyanin components are anthocyanidin 3*-O-*glucoside-2”*-O-*xylosylderivatives (Table S1). This phenomenon implies that, although the first glycosylation step of anthocyanins is similar for the production of anthocyanidin 3-*O*-glucosides and is catalyzed by UDP-glucose:flavonoid 3*-O-*glucosyltransferases, further modifications of anthocyanidin 3*-O-*glucosides diverge based on the utilization of different sugar donors, i.e. glucose in sweetpotato and xylose in *Arabidopsis*. In *Arabidopsis*, glycosyltransferase UGT79B1 (At3GGT) catalyzes the conversion of UDP-xylose and cyanidin 3*-O-*glucoside into cyanidin 3*-O-*glucoside-2”*-O-*xyloside (Tohge et al., 2005; Saito et al., 2013). In sweetpotato, a novel glucosyltransferase, UDP-glucose:anthocyanidin 3*-O-*glucoside-2*-O-*glucosyltransferase (Ib3GGT), was predicted to participate in further glycosylation.

### Cloning and phylogenetic characterization of Ib3GGT

The full-length *Ib3GGT* CDS sequence (GenBank accession number EF108571) was identified from a sweetpotato cDNA library by comparison with the *At3GGT* sequence. The 1380-bp *Ib3GGT* gene harbors an open-reading frame encoding 459 amino acids (aa) with a calculated molecular mass of 50.87 kDa and an isoelectric point 6.537. Further sequence analysis of Ib3GGT showed that its amino acid sequence shared the common domain of PSPG box (334–377 aa, Fig. 1A) with other UF3GGTs in the C-terminal region (Osmani et al., 2009). In addition, although the sugar donor specificity was reported to be partially determined by the last amino acid residue of the PSPG box, i.e., glutamine (Gln) for UDP-glucose and histidine (His) for UDP-galactose (Kubo et al., 2004), the last residue of PSPG in Ib3GGT (at 377 aa), At3GGTF (UDP-glucose:flavonol 3*-O-*glucoside-2”*-O-*glucosyltransferase), At3GGT, and Ip3GGT are Gln that is conserved among these glycosyltransferases. This phenomenon indicated that other amino acid residues in their sequences might contribute towards sugar donor specificity, which necessitates further elucidation.

**Figure 1.**
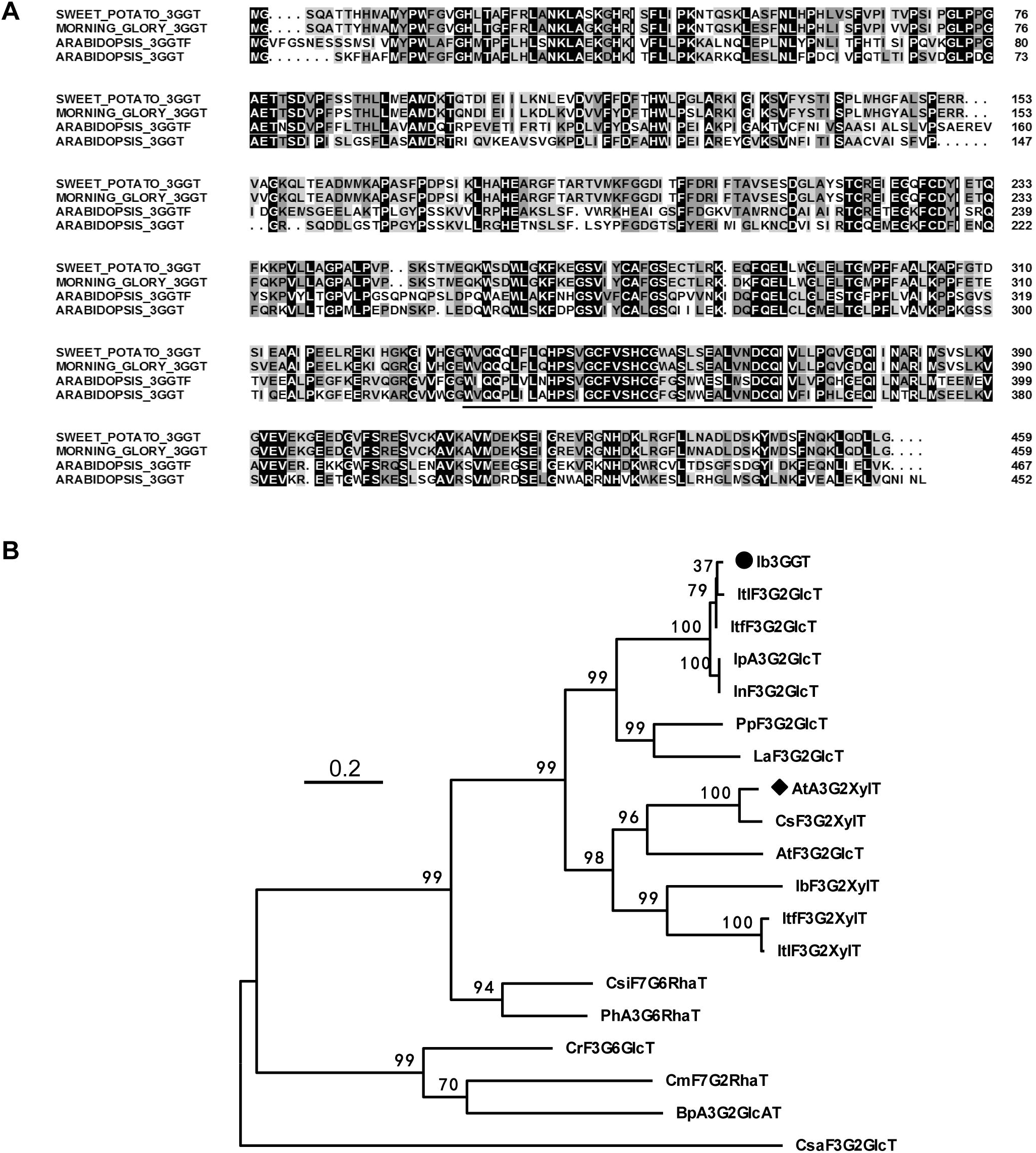
Alignment of amino acid sequences and phylogenetic tree of flavonoid glycosyltransferases. **(A)** Multiple alignments of amino acid sequences of sweetpotato Ib3GGT, morning glory Ip3GGT, and *Arabidopsis* At3GGT. The underlined nucleotides represent the putative C-terminal UDP binding motif for glycosyltransferases (PSPG). **(B)** Non-rooted molecular phylogenetic tree of flavonoid glycosyltransferases from selected plant UDP-glycosyltransferases. All amino acids were aligned using CLUSTALW. Bootstrap values from 100 retrials are indicated at each branch. The scale shows 0.2 amino acid substitution per site. The GenBank accession numbers or genome sequence codes for the sequences are shown in parentheses: AtA3G2”XylT (NP_200217); AtF3G2”GlcT (NP_200212); BpA3G2”GlcAT (AB190262); CmF7G2”RhaT (AY048882); CrF3G6”GlcT (BAH80312); CsF3G2”XylT(XP_018450414); CsaF3G2”GlcT(CCG85331); CsiF7G6”RhaT (NP_001275829); Ib3GGT (ABL74480); IbF3G2”XylT (XP_019151635); IpA3G2”GlcT (AB192315); InF3G2”GlcT (XP_019194233); ItF3G2”GlcT (itf02g12970.t1); ItF3G2”GlcT (itb02g08330.t1); ItF3G2”XylT (itb03g28310.t1); ItF3G2”XylT (itf03g22690.t2); LaF3G2”GlcT(XP_019424989); PhA3G6”RhaT (CAA81057); PpF3G2”Glc (XP_007213494). Abbreviations for species: Ac, *Actinidia chinensis*; At, *Arabidopsis thaliana*; Bp, *Bellis perennis*; Cm, *Citrus maxima*; Cr, *Catharanthus roseus*; Cs, *Camelina sativa*; Csa, *Crocus sativus*; Csi, *Citrus sinensis*; Ib, *Ipomoea batatas*; In, *Ipomoea nil*; Ip, *Ipomoea purpurea*; Itf, *Ipomoea trifida*; Itl, *Ipomoea triloba*; La, *Lupinus angustifolius*; Ph, *Petunia hybrida*; Pp, *Prunus persica*.

Phylogenetic analysis showed that Ib3GGT belongs to a cluster of typical further glycosyltransferases, and is most closely related to Ip3GGT of *Ipomoea purpurea* (Morita et al., 2005), showing 94.3% identity (Fig. 1B). Ib3GGT is also homologous to At3GGT and At3GGTF with 45.7% and 45.6% identity, respectively.

### Ib3GGT is an enzyme that catalyzes the glycosylation of anthocyanidin 3-*O*-glucoside into anthocyanidin 3-*O*-sophoroside

To further examine the function of Ib3GGT *in vitro*, recombinant His-tag fusion Ib3GGT and At3GGT proteins expressed in the yeast expression vector pYES2 (Invitrogen, USA), were used for assessing the enzymatic activity. The specificity of Ib3GGT was examined using different sugar acceptors and donors. The recombinant Ib3GGT protein only catalyzed the conversion of cyanidin 3*-O-*glucoside into cyanidin 3*-O-*sophoroside using UDP-glucose as a sugar donor (Fig. 2A). Other glucosyl acceptors such as cyanidin, cyanidin 3,5*-O-*diglucoside, and flavonol 3*-O-*glucoside could not serve as substrates, and hence, no product was detected (Fig. 2B, 2C, 2D), similar to the negative control (empty vector) (Fig. 2E). In addition, the Ib3GGT protein could use peonidin 3*-O-*glucoside as the glycosyl acceptor to form peonidin 3*-O-*sophoroside (Fig. 2F). These findings indicated that Ib3GGT used anthocyanidin 3*-O-*glucoside as the glycosyl acceptor.

**Figure 2.**
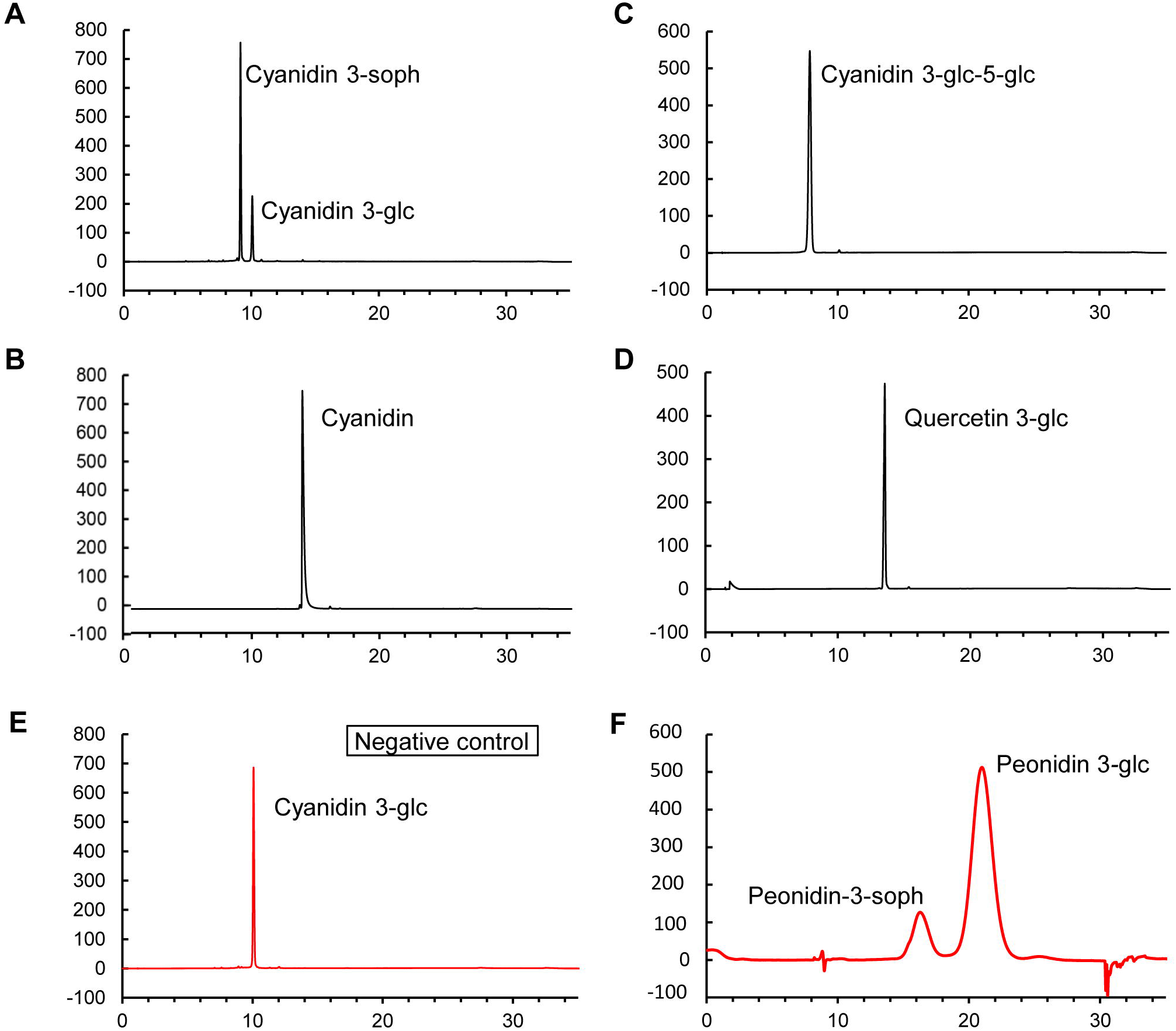
Functional assays of Ib3GGT recombinant protein using UDP-glucose and different acceptor substrates by HPLC. **(A)** Cyanidin 3-*O*-glucoside as acceptor substrate; **(B)** Cyanidin as acceptor substrate. **(C)** Cyanidin 3,5*-O-*diglucoside as acceptor substrate. **(D)** Quercetin 3*-O-*glucoside as acceptor substrate. **(E)** Cyanidin 3*-O-*glucoside as acceptor substrate without Ib3GGT protein treatment. **(F)** Peonidin 3*-O-*glucoside as acceptor substrate.

Ib3GGT specificity was also confirmed using four different UDP-sugars (Table 1). No detectable or predominant UGT activity was detected by UDP-sugars except UDP-glucose, indicating that Ib3GGT is highly specific to UDP-glucose. The weak utilization of UDP-xylose to produce cyanidin 3*-O-*glucoside-2”*-O-*xyloside indicated the low affinity to this substrate. In contrast, the At3GGT protein was capable of using only UDP-xylose (not for UDP-glucose) as the sugar donor to catalyze cyanidin 3*-O-*glucoside into cyanidin 3*-O-*glucoside-2”*-O-*xyloside (Table 1, Yonekura-Sakakibara et al., 2012), showing the divergence in the specificity of sugar donors by the two UF3GGTs from different species.

**Table 1:**
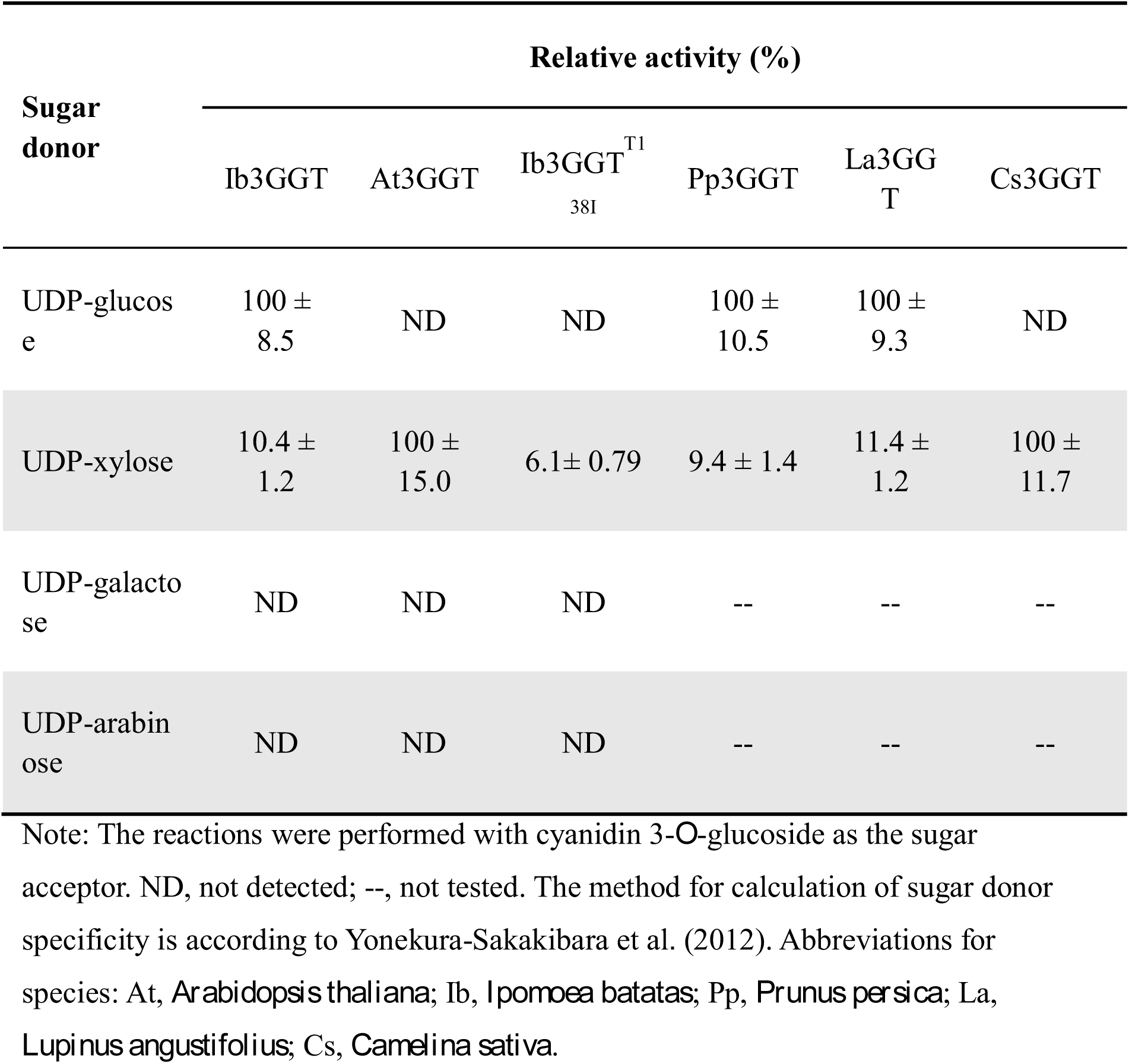
Sugar substrate specificity of different 3GGT proteins

### Thr-138 of Ib3GGT contributes to sugar donor preference

To further identify the key amino acid residue of Ib3GGT responsible for sugar donor recognition, docking experiments were performed based on the 3D structures of over 10 different glycosyltransferases enzymes from various plants (Brazier-Hicks et al., 2007; Hiromoto et al., 2006; Hiromoto et al.,2013; Modolo et al., 2009; Offen et al., 2006; Shao et al., 2005; Wetterhorn et al., 2016). The overall structures of these glycosyltransferases share a similar folding topology: two Rossmann-like domains formed a cleft which contains two substrates binding sites and one functional conserved histidine residues located between these sites (Supplementary Fig.S1). By using DALI services 8 (Holm and Laakso, 2016), more UGT homologous structures were analyzed and the root mean square deviations (RMSDs) between each other ranged from 1.1 to 2.5 Å over the core structure region (Supplementary Fig. S2). Although the sugar acceptor ligands in these structures are diverse, the sugar donors (most of them are UDP-glucose) are similar and share a group of conserved residues in their binding pockets (Supplementary Fig. S2).

Two different methods, SWISS-MODEL (Biasini et al., 2014) and I-TASSER Suite (Yang et al., 2015), were used for Ib3GGT structure modeling building. To qualify the modeling results, the overall structures of Ib3GGT with the two service models were compared with proteins 2VG8 and 2VCH; then the corresponding residues in the modeling structures were also checked. Both sets of results were similar, especially in the ligand binding sites (Supplementary Fig. S3). The RMSD between two Ib3GGT modeling structures is 1.72Å and two At3GGT modeling structures is 1.92Å, showing similar structure folding of the two services. Thus, the results of SWISS-MODEL service were used to compare the Ib3GGT/At3GGT modeling structures with an *Arabidopsis thaliana O*-glucosyltransferase (PDB ID: 2VCE, Brazier-Hicks et al., 2007) by superposing all three structures together. The UDP-glucose in the template structure (2VCE) was also well bound in *Ib*3GGT/*At*3GGT modeling structures. In Ib3GGT modeling structure, Thr22/Ser276/Glu360/Gln337/Gln338/Trp334 form a binding pocket and interact with the uridine group of UDP-glucose (Fig. 3A, Supplementary Fig.S1B); Glu277/His352/Ser357 show tightly interactions with diphosphate group (Supplementary Fig.S1C). All these residues are extremely conserved in UGTs (Hsu et al., 2018; Thompson et al., 2017).

**Figure 3.**
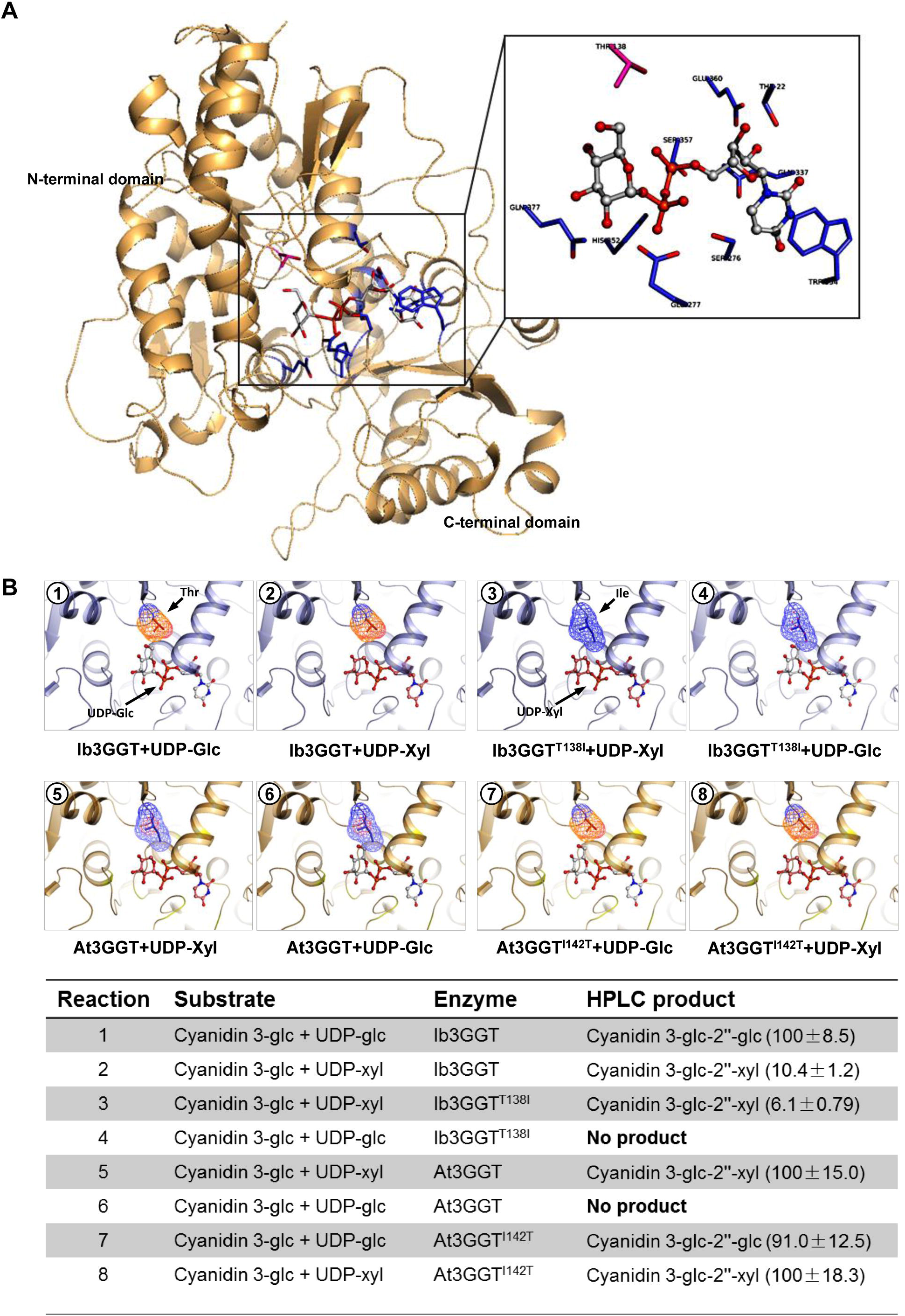
Three-dimensional modeling of Ib3GGT and At3GGT interacting with a sugar donor and a glycone acceptor. **(A)** Active center of Ib3GGT showing the key amino acid residues for sugar donor and acceptor positions. **(B)** Docking illustration of sugar donors and a glycone acceptor in the binding pocket of WT and mutant Ib3GGT and At3GGT. The performance of their reactions using cyanidin 3*-O-*glucoside with sugar nucleotide UDP-glucose or UDP-xylose is shown in the bottom panel. The percentage of relative enzyme activity is indicated in the parentheses.

The two modeling structures shared a common group of residues in binding the sugar donors (Glu277/Asp376/Gln377 in Ib3GGT and Gln285/Glu385/Gln386 in At3GGT). The only difference in sugar binding pocket is the Thr-138 in Ib3GGT which is equivalent to Ile-142 in At3GGT. The distance between O-6-glucose and Thr-138 is 2.7Å (Supplementary Fig. S4), which can form a tight interaction.

However, the distance in At3GGT is 1.7Å (Supplementary Fig.S4B), which is too short to bind the UDP-glucose. While replacing UDP-glucose with UDP-xylose in the same position may lead to weak interaction between the xylose group and Thr-138 in Ib3GGT (Supplementary Fig.S4C) and forms a 3.9Å hydrophobic interaction in At3GGT (Supplementary Fig. S4D). These modeling observations are consistent with our enzymatic activity assays (Table 1). Therefore, we hypothesized that the residue Thr-138 in Ib3GGT and its equivalence Ile-142 in At3GGT are the key residues for sugar donor specificity in purple sweetpotato and Arabidopsis, respectively.

To further attest to our hypothesis, firstly the corresponding site-directed mutants, namely Ib3GGT^T138I^ (Thr-138 changed to Ile-138) and At3GGT^I142T^ (Ile-142 changed to Thr-142), were constructed. Their enzyme activity showed that both the intact protein and Ib3GGT^T138I^ could catalyze UDP-xylose to cyanidin 3*-O-*glucoside-2”*-O-*xyloside. However, the Ib3GGT^T138I^ mutant failed to use UDP-glucose (Fig. 3B). On the other hand, At3GGT^I142T^ could not only primarily catalyze UDP-xylose to produce cyanidin 3*-O-*glucoside-2”*-O-*xyloside but also use UDP-glucose to synthesize cyanidin 3*-O-*sophorosides. These findings confirmed that T138 is a key residue for sugar (glucose/xylose) recognition in Ib3GGT.

To check whether the Thr-138 is a key residue for other sugar recognition, 3D models generated for UDP-galactose and UDP-arabinose were compared in the same position with UDP-glucose and UDP-xylose. The only difference between UDP-galactose and UDP-glucose is the direction of O_4_H moiety which changes the distance between O_4_H moiety with the main chain N from 3.1Å to 4.93Å, thus UDP-galactose should have less binding affinity than UDP-glucose (Supplementary Fig. S5A, S5B). The UDP-arabinose also has less binding affinity than UDP-xylose as the disappearance of the interaction between O5 with His-20 (Supplementary Fig. S5C, S5D). As expected, no enzymatic activities were detected for Ib3GGT or Ib3GGT ^T138I^ using UDP-galactose and UDP-arabinose as sugar donors (Table 1).

To verify whether other species have the same mechanism of sugar donor selectivity, two 3GGT proteins containing Thr-138 residue from *Prunus persica* (Pp3GGT, XP_007213494) and *Lupinus angustifolius* (La3GGT, XP_019424989) and one containing Ile-138 from *Camelina sativa* (Cs3GGT, XP_018450414) were cloned (Supplementary Fig. S6). Both Pp3GGT and La3GGT prefer UDP-glucose rather than UDP-xylose as sugar donor (Table 1). Their weak utilization of UDP-xylose indicated the low affinity to this substrate. On the other hand, the Cs3GGT protein was capable of using only UDP-xylose as the sugar donor (Table 1). Therefore, Thr-138 residue plays a key role in specificity of UDP-glucose donors by the two kinds of UF3GGTs. These results indicated that plant UGTs may share a same mechanism in sugar donor selectivity.

### *Ib3GGT* expression in *Arabidopsis* produces new anthocyanin molecules

To further validate the activity of Ib3GGT in planta, the *Ib3GGT* gene driven by the CaMV 35S promoter was overexpressed in *Arabidopsis* Col-0 and the UGT79B1 transposon insertion mutants, *ugt79b1-1* and *ugt79b1-2* (Kuromori et al., 2004; Ito et al., 2005). More than 10 independent transgenic plant lines were produced for each transformation event and their T3 homozygous lines (Fig. 4A). Further RT-PCR analysis confirmed the overexpression of *Ib3GGT* in these transgenic lines of Col-0, *ugt79b1-1* and *ugt79b1-2* (Fig. 4B, 4C). In the T3 homozygous Ib3GGT-OE transgenic lines, a new peak with an *m/z* value corresponding to cyanidin 3*-O-*sophoroside was detected in comparison to WT Col-0 by HPLC-electrospray ionization (ESI)-tandem mass spectrometry (MS/MS) analysis (Fig. 4D), although the purple-color phenotype and anthocyanin content at the cotyledon-stage seedling was indistinguishable (Fig. 4A, 4E. Moreover, the seedlings of *ugt79b1-1* and *ugt79b1-2* lines, which lacked the purple coloration as compared to the WT Nossen on 4.5% sucrose containing media, showed recovered anthocyanin accumulation when overexpressing the *Ib3GGT* gene (Fig.4A, bottom panel, Supplementary Fig. S7). The comparison of anthocyanin profiles among WT, *ugt79b1-1,* and *ugt79b1-1* overexpressing *Ib3GGT* showed the production of cyanidin 3*-O-*sophoroside in transgenic lines (Fig. 4D; Supplementary Fig. S7). These results confirmed that Ib3GGT could specifically catalyze the conversion of cyanidin 3*-O-*glucoside to cyanidin 3*-O-*sophoroside in *Arabidopsis*, a biological process absent in this plant.

**Figure 4.**
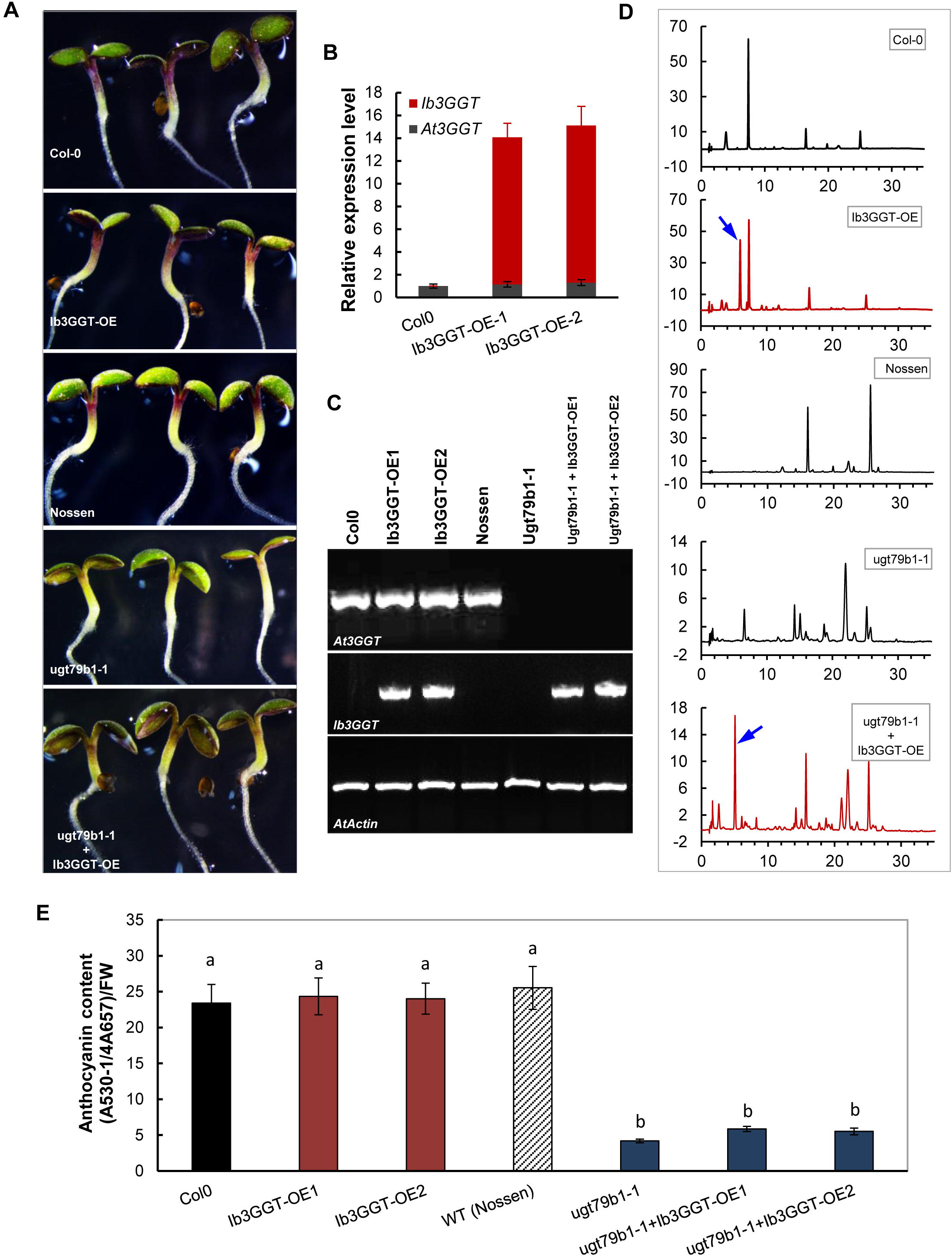
Anthocyanin characterization of transgenic *Arabidopsis* plants overexpressing *Ib3GGT* gene. **(A)** Anthocyanin pigmentation in seedlings of WTs (Col0 and Nossen), *Ib3GGT*-overexpressing Col0 lines (Ib3GGT-OE), *ugt79b1* mutant (ugt79b1-1), and Ib3GGT-overexpressing *ugt79b1* line (ugt79b1-1+Ib3GGT-OE). **(B)** The expression of *At3GGT* and *Ib3GGT* in the WT Col0 and *Ib3GGT*-overexpressing Col0 lines (Ib3GGT-OE1 and Ib3GGT-OE2) by real-time RT-PCR analysis. **(C)** RT-PCR detection of *At3GGT* and *Ib3GGT* expression in the WTs (Col0 and Nossen), two independent *Ib3GGT*-overexpressing Col0 lines (Ib3GGT-OE1 and Ib3GGT-OE2), *ugt79b1* mutant, and two *Ib3GGT*-overexpressing *ugt79b1* lines (ugt79b1-1+Ib3GGT-OE1 and ugt79b1-1+Ib3GGT-OE2). Different letters indicate significant differences (one-way ANOVA, *P* < 0.05). **(D)** Anthocyanin component profiles by HPLC/PDA/MS in the seedlings of WTs (Col0 and Nossen), *ugt79b1* mutant, and Ib3GGT-overexpressing *ugt79b1* line (ugt79b1-1+Ib3GGT-OE). Blue arrows indicate the new peaks of cyanidin 3*-O-*sophoroside. **(E)** Anthocyanin content in the WTs (Col0 and Nossen), two independent *Ib3GGT*-overexpressing Col0 lines (Ib3GGT-OE1 and Ib3GGT-OE2), *ugt79b1* mutant, and two *Ib3GGT*-overexpressing *ugt79b1* line (ugt79b1-1+Ib3GGT-OE1 and ugt79b1-1+Ib3GGT-OE2).

### *Ib3GGT* expression is associated with anthocyanin accumulation and organ development and is regulated by IbMYB1

Anthocyanin accumulation in Ayamurasaki plants showed an organ-dependent pattern. The immature leaves and mature storage roots contained maximum levels of anthocyanins (Fig. 5A); while mature leaves and fibrous roots had the least amounts. Among leaves, Lf1 reached a concentration of 0.6324 mg/g, approximately 7-fold that of Lf5. The *Ib3GGT* expression analyzed by real-time PCR in different organs also showed a similar pattern - high expression was found in immature leaves as well as developing and mature storage roots (Fig. 5A). *Ib3GGT* was expressed abundantly in developing storage roots (Dt, Fig. 5A) as compared to mature roots (Mt), which accumulated 30% more anthocyanins (0.4276 mg/g, Fig. 5A). Overall, *Ib3GGT* expression was associated with anthocyanin accumulation in different organs of Ayamurasaki plants.

**Figure 5.**
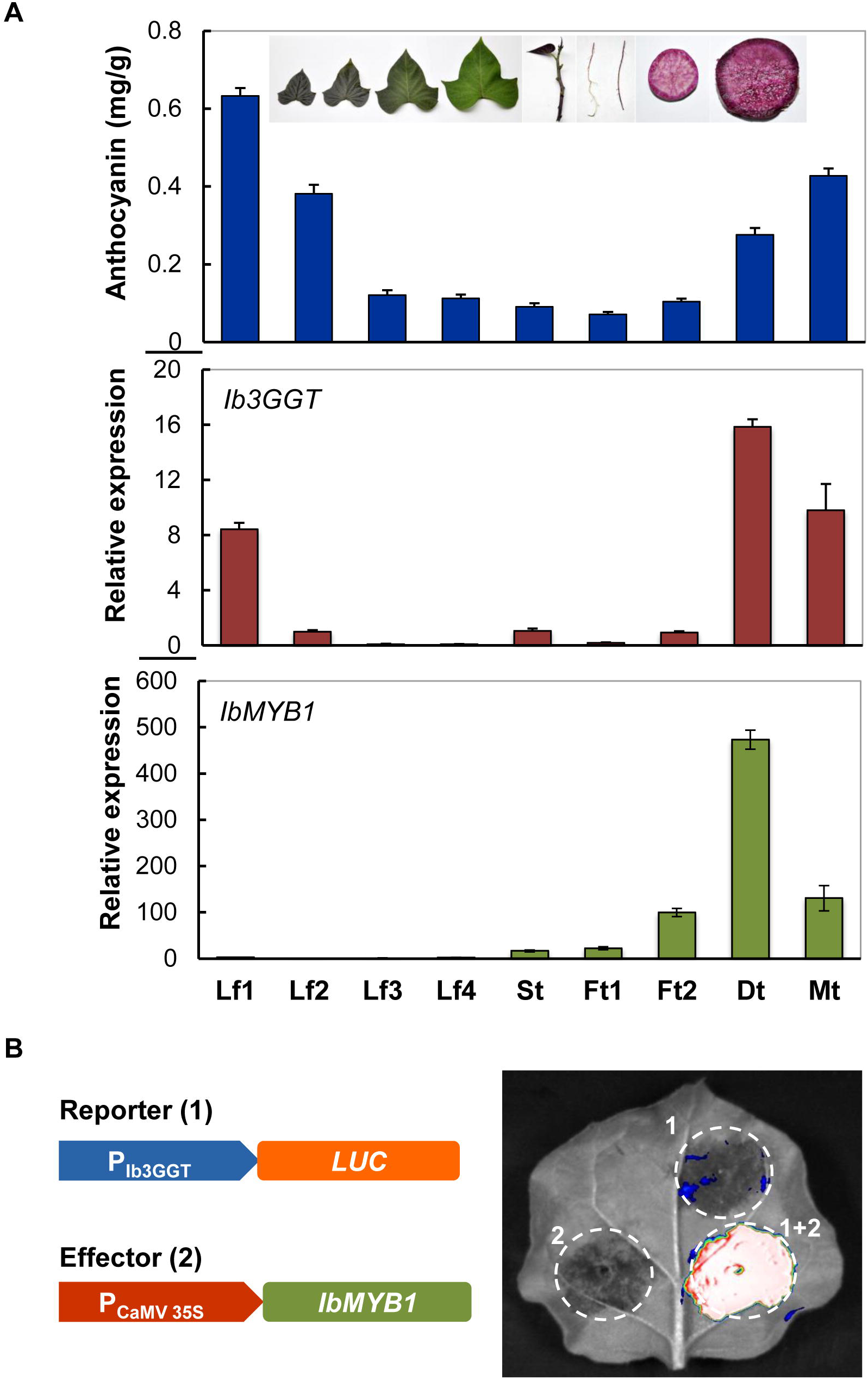
Correlation of anthocyanin accumulation and gene expression in various organs of sweetpotato cv. Ayamurasaki. **(A)** Profiles of anthocyanin accumulation, and *Ib3GGT* and *IbMYB1* transcript levels as detected by qRT-PCR in different organs. Lf1, Lf2, Lf3, and Lf4 represent leaves of different developmental stages: St, stem; Ft1, white fibrous root; Ft2, red fibrous root; Dt, developing root; Mt, mature root. Values are mean ± SD (n = 6). **(B)** Luciferase assay of *Ib3GGT* promoter activity (reporter) regulated by IbMBY1 (effector) in agroinfiltrated tobacco leaves.

The transcription factor IbMYB1 predominantly regulates the anthocyanin biosynthesis in purple sweetpotato (Mano et al., 2007), and its expression in Ayamurasaki plants was associated with anthocyanin accumulation. Therefore, *Ib3GGT* could potentially be a target gene of IbMYB1 that regulates the expression of downstream genes by binding the G-box element (CACGTG) in their promoters (Mano et al., 2007). Nonetheless, the promoter region of *Ib3GGT* showed a G-box element at position 992 as analyzed by the PlantCARE software (Supplementary Fig. S8). To confirm that *Ib3GGT* expression was stimulated by IbMYB1, a luciferase gene reporter driven by a 2000 bp promoter of *Ib3GGT* was assayed for luciferase activity in tobacco leaves after co-agroinfiltration with the effector, which harbors the *CaMV 35S::IbMYB1* expression cassette. Interestingly, a strong luciferase activity was detected. Reporter only or effector with a corresponding empty vector failed to detect the luminescent signals (Fig. 5B). These findings indicated that IbMYB1 regulates the *Ib3GGT* expression in sweetpotato Ayamurasaki plants.

### Regulation of *Ib3GGT* expression in sweetpotato Ayamurasaki alters the anthocyanin content but not the overall component profile

To further elucidate the role of Ib3GGT in sweetpotato, *Ib3GGT*-overexpressing (OE-Ib3GGT) or -RNAi (RNAi-Ib3GGT) transgenic Ayamurasaki plants were analyzed. Multiple independent transgenic plant lines were produced and propagated in the greenhouse. Compared to the WT, RNAi-Ib3GGT lines showed reduced anthocyanin levels in the leaves of pot-grown plants, whereas OE-Ib3GGT plants showed an increased anthocyanin accumulation in the top leaves (Fig. 6A, 6B). The expression of *Ib3GGT* was down-regulated in the RNAi-Ib3GGT lines and up-regulated in the OE-Ib3GGT lines by real-time PCR analyses (Fig. 6C). Anthocyanin level in the third leaf was reduced to 28.5% of WT in the RNAi-Ib3GGT-2 and increased up to 112% of WT in the OE-Ib3GGT-2 (Fig. 6A, 6B, 6D). The changes in the anthocyanin levels were correlated to *Ib3GGT* expression in these plants (Fig. 6C, 6D). Nevertheless, the overall profile of anthocyanins did not alter in these plants (Fig. 6E), indicating that Ib3GGT is involved in an early stage of anthocyanin modifications. Furthermore, the auto-fluorescence assayed in leaf epidermal cells showed a dramatic reduction in the fluorescent intensity in RNAi-Ib3GGT lines, while WT and OE-Ib3GGT transgenic plants displayed strong signals (Fig. 6F). Similar trends of altered anthocyanin accumulation were also observed in the field-grown corresponding plants (Supplementary Fig. S9).

**Figure 6.**
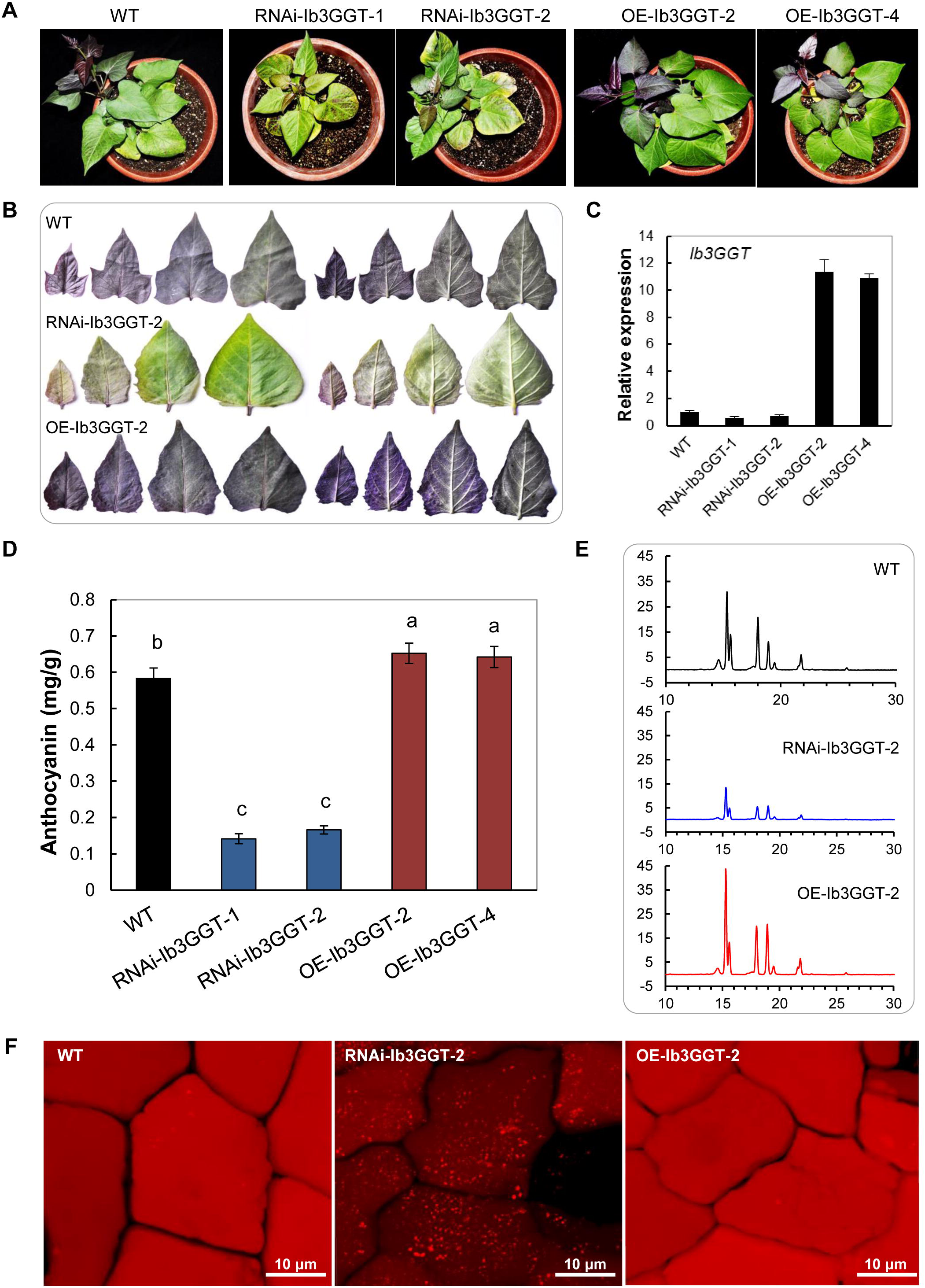
Anthocyanin characterization of wildtype and *Ib3GGT* transgenic sweetpotato plants. **(A)** Pot-grown plant phenotypes. WT, wild type; OE-Ib3GGT line, transgenic plants overexpressing *Ib3GGT*; RNAi-Ib3GGT line, *Ib3GGT* RNAi transgenic plants. **(B)** Anthocyanin pigmentation in top leaves of WT, RNAi-Ib3GGT-2, and OE-Ib3GGT-2 plants. Both the adaxial (left) and abaxial (right) leaf surfaces are shown. **(C)** Relative transcription levels of native *Ib3GGT* and the *Ib3GGT* transgene in WT and transgenic lines assessed by qRT-PCR. **(D)** Anthocyanin content in WT, RNAi-Ib3GGT, and OE-Ib3GGT plant lines. Different letters indicate significant differences (one-way ANOVA, *P* < 0.05). **(E)** Component profiles of anthocyanins in WT, RNAi-Ib3GGT-2, and OE-Ib3GGT-2 plants, as assessed by HPLC. **(F)** Anthocyanin autofluorescence in leaf epidermal cells of RNAi-Ib3GGT-2 and OE-Ib3GGT-2 plants.

### Ib3GGT functions in the cytosol

Anthocyanins have been suggested to be synthesized on the outer surface of the ER. To determine the location of the glycosylation of anthocyanins by Ib3GGT, the putative transit peptides were predicted *in silico* by Signal IP3.0, but none was found in the full Ib3GGT protein sequence. To test whether Ib3GGT is associated with ER, the N- and C-terminal fusions of Ib3GGT to eGFP regulated by the CaMV 35S promoter, together with an ER-marker or a soluble mRFP, were transiently expressed in *Nicotiana benthamiana* leaves (Fig. 7). Both, the N- and C-terminal Ib3GGT fusion proteins were found to localize in the cytosol, similar to the soluble mRFP (Fig. 7C, 7D). However, some signal of the eGFP-Ib3GGT fusions was also observed in the nucleus. Additionally, when Ib3GGT-eGFP was expressed together with ER-mCherry, no co-localization was found (Fig. 7A, 7B). Thus, Ib3GGT is a soluble protein in the cytosol and not associated with the ER (Poustka et al., 2007).

**Figure 7.**
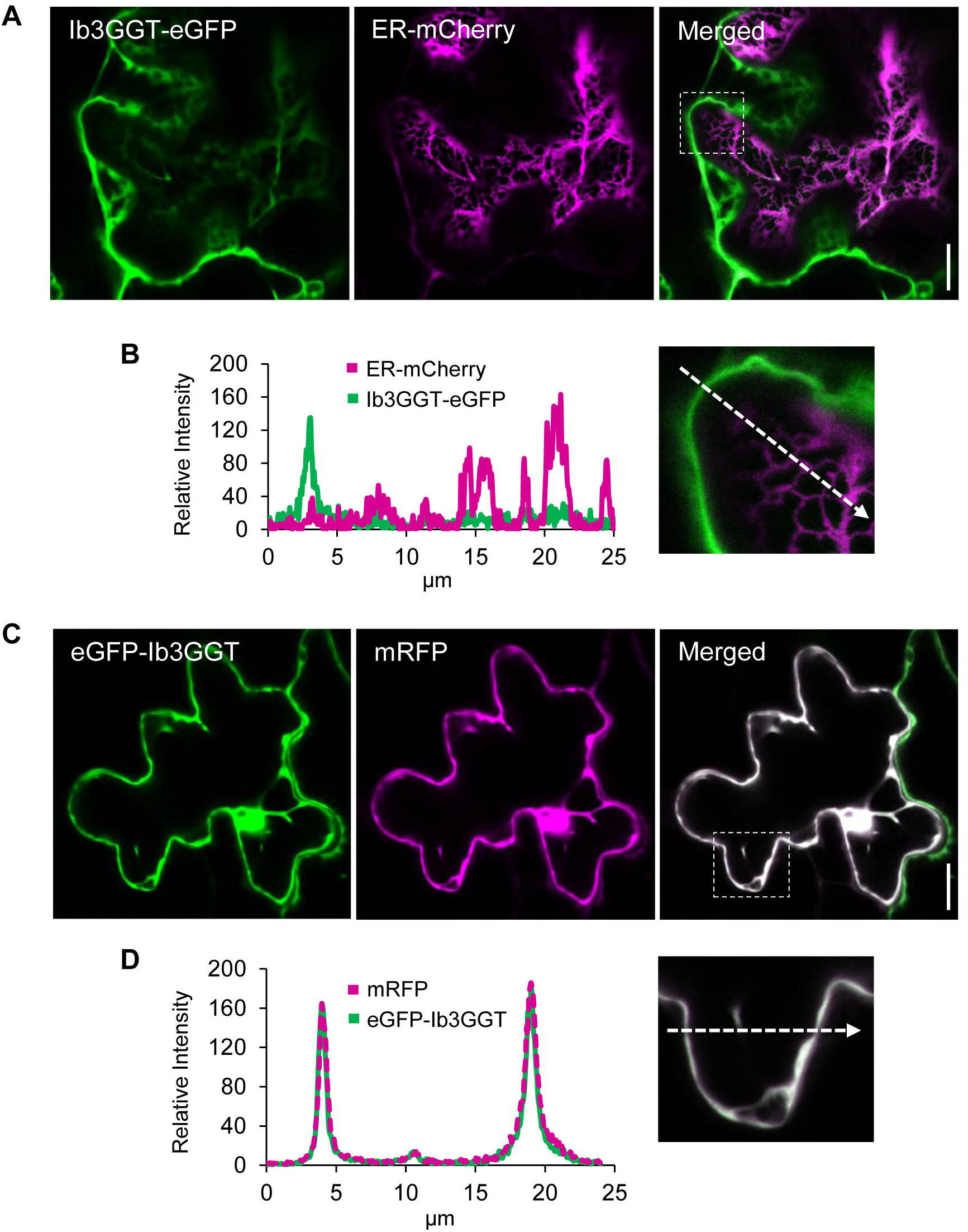
Subcellular localization of Ib3GGT after transient expression in *Nicotiana benthamiana* leaves. **(A)** Optical section through a pavement cell, co-expressing Ib3GGT-eGFP (left panel) and ER-mCherry (middle panel). GFP and mCherry signals are distinct in the merged image (right panel). The outlined region is shown at a higher magnification in (B). **(B)** Relative fluorescence intensity along the axis is marked by the dotted arrow in the right panel. **(C)** Co-expression of eGFP-Ib3GGT (left panel) and soluble mRFP (middle panel). White pixels of the merged image show an overlay of both channels (right panel). The marked region was enlarged in (D). **(D)** Intensity plot, as presented before. eGFP-Ib3GGT and soluble mRFP were co-localized in the cytoplasm. Scale bar, 15 µm.

## Discussion

### Further glycosylation modifications of anthocyanins in the cytosol

Further glycosylation of anthocyanins is an essential step in their biosynthesis, accumulation, and stability (Yonekura-Sakakibara et al., 2008; Zhang et al., 2014). In purple sweetpotato, the major anthocyanins include cyanidin and peonidin 3-sophorosides as well as their acylated derivatives (Truong et al., 2009; Lee et al., 2013), indicating that further glycosylation is required for the conversion from anthocyanidin 3*-O*-glucosides into anthocyanidin 3*-O*-sophorosides. In this study, we found that Ib3GGT was responsible for the reaction using UDP-glucose as the sugar donor. Unlike sweetpotato, further glycosylation of anthocyanins is xylosylation catalyzed by UGT79B1 in *Arabidopsis* (Saito et al., 2013). Apparently, *Arabidopsis* lacks the enzymes to form cyanidin 3*-O*-sophoroside using cyanidin 3*-O*-glucosides as the substrate for further modification, as the overexpressing Ib3GGT in *Arabidopsis* only showed a peak of cyanidin 3*-O*-sophoroside without a change in other anthocyanin components. Therefore, Ib3GGT is a key player of anthocyanin for further glycosylation modifications in sweetpotato.

Further glycosylation is a critical step determining the subsequent anthocyanin modifications, such as malonylation and acylation (Yonekura-Sakakibara et al., 2008; Andersen and Jordheim, 2010). The Ib3GGT as UDP-glycosyltransferase can add a sugar residue to anthocyanidin 3-*O*-glucosides but not anthocyanidin 3,5-*O*-diglucosides (Fig. 2A, 2C). Thus, we conclude that 5GT catalyzes a glucose molecule into the 5^th^ position of C-ring that can hinder the transfer of a glucose molecule into the 2” position of anthocyanidin 3-*O*-glucosides. In certain order modification of anthocyanin, this phenomenon might be a crucial factor for providing the condition for subsequent modification in anthocyanin. Ib3GGT cannot catalyze flavonol-3-*O*-glucoside as acceptor substrate, demonstrating that Ib3GGT has substrate specificity in purple sweetpotato (Fig. 2D). However, At3GGT can catalyze flavonol-3-*O*-glucoside and anthocyanin-3-*O*-glucoside as a broad substrate in *Arabidopsis* (Saito et al., 2013).

As a primary sedative mechanism that maintains metabolic homeostasis in plants, glycosylation contributes to the diversity in synthesizing various secondary plant metabolites, thereby altering the biological functions of these metabolites (Jones and Vogt, 2001; Gachon et al., 2005). Apparently, divergence occurs among species by adaption of glycosyltransferase substrate specificity. In peach, the PpUGT79B is responsible for glycosylation by adding a rhamnoside molecule to anthocyanidin 3-*O*-glucosides forming the anthocyanidin 3-*O*-rutinoside (Cheng et al., 2014). Ib3GGT was functional in transgenic *Arabidopsis* by producing anthocyanin-3-sophoroside; while the overexpression of UGT79B1 (At3GGT) in sweetpotato did not catalyze the production of anthocyanin 3*-O-*glucoside-2”*-O-*xylose (Supplementary Fig. S10). These findings indicated that anthocyanin glycosylation in sweetpotato diverges from that of *Arabidopsis* towards the specific sugar acceptor. Interestingly, the further glycosylation of anthocyanin occurs in the cytosol, not like other UGTs that are mainly ER membrane-bound enzymes (Poustka et al., 2007; Zhao, 2015).

### Key amino acids in UGTs affect both sugar donor preference and regioselectivity

The phylogenetic comparisons of flavonoid GGTs suggested that the potentially conserved amino acid residues are involved in further substrate-selectivity. Four amino acid residues (Trp-334/Gln-337/Glu-360/His-352 in Ib3GGT) are generally conserved across all known flavonoid 3*-O-*glycoside-2”*-O-*glycosyltransferases. The close relationship between Ib3GGT and UGT79B1 in the phylogenetic tree also indicated that the sugar donor selectivity of flavonoid GGTs was established after species differentiation (Saito et al., 2013). In sweetpotato, Ib3GGT accepts UDP-glucose as sugar donor to conjugate to anthocyanins, such as cyanidin 3-*O*-glucoside or peonidin 3-*O*-glucoside. Interestingly, *Arabidopsis* also has a UDP-glucose:flavonoid3*-O-*glucoside-2”*-O-*glucosyltransferase (At3GGTF), which preferentially uses flavonol 3*-O-*glucoside and UDP-glucose as substrates (Kubo et al., 2007). Thr-138 as the key residue for UDP-glucose recognition is also conserved in glucosyltransferases that uses UDP-glucose as sugar donor in morning glory, *Arabidopsis* (At3GGTF), *Ricinus communis*, and *Glysin max* (Supplementary Fig. S6). Corresponding to the Thr-138 residue, the Ile-142 is the residue for UDP-xylose recognition in *Arabidopsis*. Instead, the corresponding sites of Ile in *Camelina sativa*, Thr in *Tarenaya hassleriana* and *Brassica napus*, and Val in *Eucalyptus grandis* are also responsible for recognizing UDP-xylose (Supplementary Fig. S6).

### Anthocyanin glycosyltransferases are regulated by transcription factors (TFs)

Anthocyanin biosynthesis is a finely regulated system involving multiple TFs associated with plant development (Pireyre and Burow, 2015; Xu et al., 2015). For example, the temporal and spatial regulation of anthocyanin production in flowers mediated by TFs, R2R3-MYB, basic Helix-Loop-Helix (bHLH), or WD40 type (reviewed in Davies et al., 2012) brings the colorful variation in the world. Hitherto, the only well-characterized TF in sweetpotato is R2R3-MYB type IbMYB1, which controls anthocyanin biosynthesis specifically in tuberous roots by inducing all the structural anthocyanin genes (Mano et al., 2007). In this study, the accumulation of anthocyanin in different organs was strongly associated with the *Ib3GGT* expression, which indicates its divergent regulation during plant development. Importantly, the activation of the *Ib3GGT* promoter by IbMYB1 confirmed that *Ib3GGT* is highly regulated by the TF in storage roots. The relatively low level of *IbMYB1* transcript in the leaves might reflect its tissue specificity. In *Arabidopsis*, it is well-documented that the R2R3-MYB TF can induce glycosyltransferases such as UGT79B1 (Tohge et al., 2005; Yonekura-Sakakibara et al., 2008; Stracke et al., 2010).

In summary, sweetpotato Ib3GGT catalyzes the anthocyanidin 3-*O*-glucosides into anthocyanidin 3-*O*-sophorosides using UDP-glucose as a sugar donor. The Thr-138 of Ib3GGT is a key residue for sugar donor selectivity in the further glycosylation that contributes to the stability and diversity of anthocyanins. The Ib3GGT glycosylation occurs in the cytosol and is regulated by IbMYB1 TF. The present study provides further insights regarding the glycosylation enzymes involved in secondary metabolism in divergence that can assist in developing a useful approach to diversifying certain flavonoids in crops.

## Supplementary data

**Fig. S1.** UDP-glucose binding sites of Ib3GGT. (A) Chemical structure of UDP-glucose. (B,C) Close-up views of the interactions of uridine moiety (B), and diphosphate moiety. (D) Overall structure of Ib3GGT modeling structure. UDP-glucose is showed with ball-and-stick model and the protein structure is showed with cartoon; the helix/sheet/loops are showed in cyan/red/magenta, respectively.

**Fig. S2.** Structure alignment of UGT homologs. (A) All structures are showed with ribbon and superposed by Coot (Emsley et al., 2010). The PDB numbers of these structures are 2ACW, 2C1Z, 2PQ6, 2VG8, 3HBF, 3WC4, 5GL5, 5NLM, 5TMB and 5U6M. (B) Ligands in superposed structures are showed with sticks.

**Fig. S3.** Sequence alignment of Ib3GGT with 2VG8 and 2VCH proteins using SWISS-MODEL and I-TASSER.

**Fig. S4.** Sugar donor binding sites in Ib3GGT and At3GGT. (A, B, C, D) Close-up views of the interactions of UDP-glucose with Ib3GGT (A), UDP-glucose with At3GGT (B), UDP-xylose with Ib3GGT (C), and UDP-xylose with At3GGT (D). UDP-glucose and UDP-xylose are showed with stick-and-ball in magenta and red, respectively; side chain residues are showed with stick; the hydrogen bonds are indicated by dashed lines.

**Fig. S5.** Difference in binding affinity of sugar analogies. (A, B, C, D) Ball-and-stick model of UDP-galactose (A), UDP-glucose (B), UDP-arabinose (C), and UDP-xylose (D). They are showed in grey, magenta, pink, and yellow, respectively. Chemical structure of galactose, glucose, arabinose, and xylose are showed in the corresponding positions.

**Fig. S6.** Amino acid sequence comparison of GGT analogies with Ib3GGT. The Thr-138 site is boxed.

**Fig. S7.** Anthocyanin pigmentation and component profiles in seedlings of the *ugt79b1-2* mutant and *Ib3GGT*-overexpressing *ugt79b1-2* transgenic line (ugt19b1-2+Ib3GGT-OE).

**Fig. S8.** *Ib3GGT* promoter sequence showing the two G-box sites.

**Fig. S9.** Leaf and root phenotypes of field-grown wild-type (WT), RNAi-Ib3GGT-2 and OE-Ib3GGT-2 plant lines.

**Fig. S10.** Analysis of anthocyanin compounds in wild-type (WT) and

*At3GGT*-overexpressing (OE-At3GGT) sweetpotato by HPLC-MS.

**Table S1:** Anthocyanin compounds in sweetpotato (Tian et al., 2005) and *Arabidopsis* (Tohge et al., 2005)

**Table S2:** List of primers for gene expression analysis in *Arabidopsis* and sweetpotato plant lines.

## Acknowledgments

This work was supported by the grants from the National Natural Science Foundation of China (31501357, 31771854), the Collaborative Innovation Action – Agricultural

Science and Technology Innovation Program of Chinese Academy of Agricultural Sciences (CAAS-XTCX2016009), and XDPB0402 of CAS. We thank Prof. Cathie Martin from JIC for providing helpful suggestions and corrections for the manuscript. We also thank our colleagues Prof. Peng Zhang for protein structure analysis, Miss Yuanhong Shan for HPLC-MS-MS analysis and Mr. XiaoshuGao for confocal microscopy.

## Reference

Andersen ØM, Jordheim M. 2010. Anthocyanins. In: Encyclopedia of Life Sciences (ELS). Chichester: John Wiley and Sons Ltd.

Biasini M, Bienert S, Waterhouse A, Arnold K, Studer G, Schmidt T, Kiefer F, Gallo CT, Bertoni M, Bordoli L. 2014. SWISS-MODEL: modelling protein tertiary and quaternary structure using evolutionary information. Nucleic Acids Research 42, W252-W258.

Brazier-Hicks M, Offen WA, Gershater MC, Revett TJ, Lim EK, Bowles DJ, Davies G J, Edwards R. 2007. Characterization and engineering of the bifunctional N- and O-glucosyltransferase involved in xenobiotic metabolism in plants. Proceedings of the National Academy of Sciences, USA 104, 20238–20243.

Breton C, Šnajdrová L, Jeanneau C, Koča J, Imberty A. 2006. Structures and mechanisms of glycosyltransferases. Glycobiology 16, 29R–37R.

Caputi L, Malnoy M, Goremykin V, Nikiforova S, Martens S. 2012. A genome-wide phylogenetic reconstruction of family 1 UDP-glycosyltransferases revealed the expansion of the family during the adaptation of plants to life on land. Plant Journal 69, 1030–1042.

Cheng J, Wei G, Zhou H, Gu C, Vimolmangkang S, Liao L, Han Y. 2014. Unraveling the mechanism underlying the glycosylation and methylation of anthocyanins in peach. Plant Physiology 166, 1044–1058.

Claudia C, Philippe R, Edith F, Vincent B, Yumei Z, Sharyn EP, Juan J R, Martin Y, Christophe D. 2017. The class III peroxidase PRX17 is a direct target of the MADS-box transcription factor AGAMOUS-LIKE15 (AGL15) and participates in lignified tissue formation. New Phytologist 213, 250–263.

Clough SJ, Bent AF. 1998. Floral dip: a simplified method for Agrobacterium-mediated transformation of Arabidopsis thaliana. Plant Journal 16, 735–743.

Davies KM, Albert NW, Schwinn KE. 2012. From landing lights to mimicry: the molecular regulation of flower colouration and mechanisms for pigmentation patterning. Funct. Plant Biology 39, 619–638.

de Pascual-Teresa S, Sanchez-Ballesta MT. 2008. Anthocyanins: from plant to health. Phytochemistry Reviews 7, 281–299.

Emsley P, Lohkamp B, Scott WG, Cowtan K. 2010. Features and development of Coot. Acta Crystallographica Section D: Structural Biology 66, 486–501.

Fedoroff NV, Furtek DB, Nelson OE. 1984. Cloning of the bronze locus in maize by a simple and generalizable procedure using the transposable controlling element Activator (Ac). Proceedings of the National Academy of Sciences, USA 81, 3825–3829.

Gachon CM, Langlois-Meurinne M, Saindrenan P. 2005. Plant secondary metabolism glycosyltransferases: the emerging functional analysis. Trends in Plant Science 10, 542–549.

Glover BJ, Martin C. 2012. Anthocyanins. Current Biology 22, R147–R150.

Griesser M, Hoffmann T, Bellido ML, Rosati C, Fink B, Kurtzer R, Aharoni A, Munoz-Blanco J, Schwab W. 2008a. Redirection of flavonoid biosynthesis through the down-regulation of an anthocyanidin glucosyltransferase in ripening strawberry fruit. Plant Physiology 146, 1528–1539.

Griesser M, Vitzthum F, Fink B, Bellido ML, Raasch C, Munoz-Blanco J, Schwab W. 2008b. Multi-substrate flavonol O-glucosyltransferases from strawberry (Fragaria× ananassa) achene and receptacle. Journal of Experimental Botany 59, 2611–2625.

He J, Giusti MM. 2010. Anthocyanins: natural colorants with health-promoting properties. Annual Review of Food Science and Technology 1, 163–187.

He ZH, Cheeseman I, He D, Kohorn BD. 1999. A cluster of five cell wall-associated receptor kinase genes, Wak1-5, are expressed in specific organs of Arabidopsis. Plant Molecular Biology 39, 1189–1196.

Hiromoto T, Honjo E, Noda N, Tamada T, Kazuma K, Suzuki M, Blaber M, Kuroki R. 2015. Structural basis for acceptor-substrate recognition of UDP-glucose:anthocyanidin 3-O-glucosyltransferase from Clitoria ternatea. Protein Science 24, 395–407.

Hiromoto T, Honjo E, Tamada T, Noda N, Kazuma K, Suzukid M, Kurokia R. 2013. Crystal structure of UDP-glucose:anthocyanidin 3-O-glucosyltransferase from Clitoria ternatea. Journal of Synchrotron Radiation 20, 894–898.

Holm L, Laakso LM. 2016. Dali server update. Nucleic Acids Research 44, 351–355.

Hsu TM, Welner DH, Russ ZN, Cervantes B, Prathuri RL, Adams PD, Dueber JE. 2018. Employing a biochemical protecting group for a sustainable indigo dyeing strategy. Nature Chemical Biology 14, 256–261.

Ito T, Motohashi R, Kuromori T, Noutoshi Y, Seki M, Kamiya A, Mizukado S, Sakurai T, Shinozaki K. 2005. A resource of 5,814 dissociation transposon-tagged and sequence-indexed lines of Arabidopsis transposed from start loci on chromosome 5. Plant Cell Physiology 46, 1149–1153.

Jones P, Vogt T. 2001. Glycosyltransferases in secondary plant metabolism: tranquilizers and stimulant controllers. Planta 213, 164–174.

Kim HS, Kim B, Sung S, Kim M, Mok H, Chong Y, Ahn J. 2013. Engineering flavonoid glycosyltransferases for enhanced catalytic efficiency and extended sugar-donor selectivity. Planta 238, 683–693.

Kovinich N, Kayanja G, Chanoca A, Riedl K, Otegui MS, Grotewold E. 2014. Not all anthocyanins are born equal: distinct patterns induced by stress in Arabidopsis. Planta 240, 931–940.

Kroon J, Souer E, de Graaff A, Xue Y, Mol J, Koes R. 1994. Cloning and structural analysis of the anthocyanin pigmentation locus Rt of Petunia hybrida: characterization of insertion sequences in two mutant alleles. Plant Journal 5, 69–80.

Kubo A, Arai Y, Nagashima S, Yoshikawa T. 2004. Alteration of sugar donor specificities of plant glycosyltransferases by a single point mutation. Archives of Biochemistry and Biophysics 429, 198–203.

Kubo H, Nawa N, Lupsea SA. 2007. Anthocyaninless1 gene of Arabidopsis thaliana encodes a UDP-glucose:flavonoid-3-O-glucosyltransferase. Journal of Plant Research 120, 445–449.

Kuromori T, Hirayama T, Kiyosue Y, Takabe H, Mizukado S, Sakurai T, Akiyama K, Kamiya A, Ito T, Shinozaki K. 2004. A collection of 11800 single-copy Ds transposon insertion lines in Arabidopsis. Plant Journal 37, 897–905.

Lairson LL, Henrissat B, Davies GJ, Withers SG. 2008. Glycosyltransferases: structures, functions, and mechanisms. Annual Review of Biochemistry 77, 521–555.

Leuzinger K, Dent M, Hurtado J, Stahnke J, Lai H, Zhou X, Chen Q. 2013. Efficient agroinfiltration of plants for high-level transient expression of recombinant proteins. Journal of Visualized Experiments 77, 50521.

Lee MJ, Park JS, Choi DS, Jung MY. 2013. Characterization and quantitation of anthocyanins in purple-fleshed sweet potatoes cultivated in Korea by HPLC-DAD and HPLC-ESI-QTOF-MS/MS. Journal of Agricultural and Food Chemistry 61, 3148–3158.

Mano H, Ogasawara F, Sato K, Higo H, Minobe Y. 2007. Isolation of a regulatory gene of anthocyanin biosynthesis in tuberous roots of purple-fleshed sweetpotato. Plant Physiology 143, 1252–1268.

Matsuba Y, Sasaki N, Tera M, Okamura M, Abe Y, Okamoto E, Nakamura H, Funabashi H, Takatsu M, Saito M, Matsuoka H, Nagasawa K, Ozeki Y. 2010. A novel glucosylation reaction on anthocyanins catalyzed by acyl-glucose-dependent glucosyltransferase in the petals of carnation and delphinium. Plant Cell 22, 3374–3389.

Modolo LV, Li L, Pan HY, Blount JW, Dixon RA, Wang XQ. 2009. Crystal structures of glycosyltransferase UGT78G1 reveal the molecular basis for glycosylation and deglycosylation of (iso)flavonoids. Journal of Molecular Biology 392, 1292–1302.

Moglia A, Lanteri S, Comino C, Hill L, Knevitt D, Cagliero C, Rubiolo P, Bornemann S, Martin C. 2014. Dual catalytic activity of hydroxycinnamoyl-Coenzyme A quinate transferase from tomato allows it to moonlight in the synthesis of both mono-and dicaffeoylquinic acids. Plant Physiology. 166, 1777–1787.

Montefiori M, Espley RV, Stevenson D, Cooney J, Datson PM, Saiz A, Atkinson RG, Hellens RP, Allan AC. 2011. Identification and characterisation of F3GT1 and F3GGT1, two glycosyltransferases responsible for anthocyanin biosynthesis in red-fleshed kiwifruit (Actinidia chinensis). Plant Journal 65, 106–118.

Morita Y, Hoshino A, Kikuchi Y, Okuhara H, Ono E, Tanaka Y, Fukui Y, Saito N, Nitasaka E, Noguchi H. 2005. Japanese morning glory dusky mutants displaying reddish-brown or purplish-gray flowers are deficient in a novel glycosylation enzyme for anthocyanin biosynthesis, UDP-glucose:anthocyanidin 3-O-glucoside-2’’-O-glucosyltransferase, due to 4-bp insertions in the gene. Plant Journal 42, 353–363.

Nelson BK, Cai X, Nebenfuhr A. 2007. A multicolored set of in vivo organelle markers for co-localization studies in Arabidopsis and other plants. Plant Journal 51, 1126–1136.

Noguchi A, Horikawa M, Fukui Y, Fukuchi-Mizutani M, Iuchi-Okada A, Ishiguro M, Kiso Y, Nakayama T, Ono E. 2009. Local differentiation of sugar donor specificity of flavonoid glycosyltransferase in Lamiales. Plant Cell 21, 1556–1572.

Offen W, Martinez-Fleites C, Yang M, Kiat-Lim E, Davis BG, Tarling CA, Ford CM, Bowles DJ, Davies GJ. 2006. Structure of a flavonoid glucosyltransferase reveals the basis for plant natural product modification. EMBO Journal 25, 1396–1405.

Ono E, Fukuchi-Mizutani M, Nakamura N, Fukui Y, Yonekura-Sakakibara K, Yamaguchi M, Nakayama T, Tanaka T, Kusumi T, Tanaka Y. 2006. Yellow flowers generated by expression of the aurone biosynthetic pathway. Proceedings of the National Academy of Sciences, USA 103, 11075–11080.

Ono E, Homma Y, Horikawa M, Kunikane-Doi S, Imai H, Takahashi S, Kawai Y, Ishiguro M, Fukui Y, Nakayama T. 2010. Functional differentiation of the glycosyltransferases that contribute to the chemical diversity of bioactive flavonol glycosides in grapevines (Vitis vinifera). Plant Cell 22, 2856–2871.

Osmani SA, Bak S, Møller BL. 2009. Substrate specificity of plant UDP-dependent glycosyltransferases predicted from crystal structures and homology modeling. Phytochemistry 70, 325–347.

Pireyre M, Burow M. 2015. Regulation of MYB and bHLH transcription factors: a glance at the protein level. Molecular Plant 8, 378–388.

Poustka F, Irani N G, Feller A, Lu Y, Pourcel L, Frame K, Grotewold E. 2007. A trafficking pathway for anthocyanins overlaps with the endoplasmic reticulum-to-vacuole protein-sorting route in Arabidopsis and contributes to the formation of vacuolar inclusions. Plant Physiology 145, 1323–1335.

Saito K, Yonekura-Sakakibara K, Nakabayashi R, Higashi Y, Yamazaki M, Tohge T, Fernie AR. 2013. The flavonoid biosynthetic pathway in Arabidopsis: structural and genetic diversity. Plant Physiology and Biochemistry 72, 21–34.

Sasaki N, Nishizaki Y, Ozeki Y, Miyahara T. 2014. The role of acyl-glucose in anthocyanin modifications. Molecules 19, 18747–18766.

Sawada S, Suzuki H, Ichimaida F, Yamaguchi MA, Iwashita T, Fukui Y, Hemmi H, Nishino T, Nakayama T. 2005. UDP-glucuronic acid:anthocyanin glucuronosyltransferase from red daisy (Bellis perennis) flowers. Enzymology and phylogenetics of a novel glucuronosyltransferase involved in flower pigment biosynthesis. Journal of Biological Chemistry 280, 899–906.

Shao H, He XZ, Achnine L, Biount JW, Dixon RA, Wang XQ. 2005. Crystal structures of a multifunctional triterpene/flavonoid glycosyltransferase from Medicago truncatula. Plant Cell 17, 3141–3154.

Shen Y, Chen Y, Wu J, Shaner NC, Robert E, Campbell RE. 2017. Engineering of mCherry variants with long Stokes shift, red-shifted fluorescence, and low cytotoxicity. PLoS One 12(2), e0171257.

Stamatakis A. 2014. RAxML version 8: a tool for phylogenetic analysis and post–analysis of large phylogenies. Bioinformatics 30, 1312–1313.

Stracke R, Jahns O, Keck M, Tohge T, Niehaus K, Fernie AR, Weisshaar B. 2010. Analysis of PRODUCTION OF FLAVONOL GLYCOSIDES- dependent flavonol glycoside accumulation in Arabidopsis thaliana plants reveals MYB11-, MYB12- and MYB111-independent flavonol glycoside accumulation. New Phytologist 188, 985–1000.

Sun Y, Li H, Huang JR. 2012. Arabidopsis TT19 functions as a carrier to transport anthocyanin from the cytosol to tonoplasts. Molecular Plant 5, 387–400.

Sun W, Liang L, Meng X, Li Y, Gao F, Liu X, Wang L. 2016. Biochemical and molecular characterization of a flavonoid 3-O-glycosyltransferase responsible for anthocyanins and flavonolsbiosynthesis in Freesia hybrida. Front Plant Science 7, 410.

Tamura K, Stecher G, Peterson D, Filipski A, Kumar S. 2013. MEGA6: molecular evolutionary genetics analysis version 6. 0. Molecular Biology and Evolution 30, 2725–2729.

Thompson AMG, Iancu CV, Neet KE, Dean JV, Choe JY. 2017. Differences in salicylic acid glucose conjugations by UGT74F1 and UGT74F2 from Arabidopsis thaliana. Scientific Reports 7, 46629.

Tohge T, Nishiyama Y, Hirai MY, Yano M, Nakajima J, Awazuhara M, Inoue E, Takahashi H, Goodenowe DB, Kitayama M. 2005. Functional genomics by integrated analysis of metabolome and transcriptome of Arabidopsis plants over-expressing an MYB transcription factor. Plant Journal 42, 218–235.

Truong VD, Deighton N, Thompson RT, McFeeters RF, Dean LO, Pecota KV, Yencho GC. 2009. Characterization of anthocyanins and anthocyanidins in purple-fleshed sweetpotatoes by HPLC-DAD/ESI-MS/MS. Journal of Agricultural and Food Chemistry 58, 404–410.

Tian Q, Konczak I, Schwartz SJ. 2005. Probing anthocyanin profiles in purple sweet potato cell line (Ipomoea batatas L. Cv. Ayamurasaki) by high-performance liquid chromatography and electrospray ionization tandem mass spectrometry. Journal of Agricultural and Food Chemistry 53, 6503–6509.

Wang H, Fan W, Li H, Yang J, Huang J, Zhang P. 2013. Functional characterization of dihydroflavonol-4-reductase in anthocyanin biosynthesis of purple sweet potato underlies the direct evidence of anthocyanins function against abiotic stresses. PLoS One 8, e78484.

Wetterhorn KM, Newmister SA, Caniza RK, Busman K, McCormick SP, Berthiller F, Adam G, Rayment I. 2016. Crystal structure of Os79 (Os04g0206600) from Oryza sativa: A UDP-glucosyltransferase involved in the detoxification of deoxynivalenol. Biochemistry 55, 6175–6186.

Xu W, Dubos C, Lepiniec L. 2015. Transcriptional control of flavonoid biosynthesis by MYB–bHLH–WDR complexes. Trends Plant Science 20, 176–185.

Yang J, Bi HP, Fan WJ, Zhang M, Wang HX, Zhang P. 2011. Efficient embryogenic suspension culturing and rapid transformation of a range of elite genotypes of sweet potato (Ipomoea batatas [L.] Lam.). Plant Science 181, 701–711.

Yang JY, Yan RX, Roy A, Xu D, Poissonl J, Zhang Y. 2015. The I-TASSER Suite: protein structure and function prediction. Nature Methods 12, 7–8.

Yonekura-Sakakibara K, Nakayama T, Yamazaki M, Saito K. 2008. Modification and stabilization of anthocyanins. In: Anthocyanins. Springer New York, 169–190.

Yonekura-Sakakibara K, Fukushima AR, Nakabayashi K, Hanada F, Matsuda S, Sugawara E, Inoue T, Kuromori T, Ito K, Shinozaki B. 2012. Two glycosyltransferases involved in anthocyanin modification delineated by transcriptome independent component analysis in Arabidopsis thaliana. Plant Journal 69, 154–167.

Yonekura-Sakakibara K, Hanada K. 2011. An evolutionary view of functional diversity in family 1 glycosyltransferases. Plant Journal 66, 182–193.

Yonekura-Sakakibara K, Nakabayashi R, Sugawara S, Tohge T, Ito T, Koyanagi M, Saito K. 2014. A flavonoid 3-O-glucoside:2”-O-glucosyltransferase responsible for terminal modification of pollen-specific flavonols in Arabidopsis thaliana. Plant Journal 79, 769–782.

Zhang Y, Butelli E, Martin C. 2014. Engineering anthocyanin biosynthesis in plants. Current Opinion in Plant Biology 19, 81–90.

Zhao J, Huhman D, Shadle G, He XZ, Sumner LW, Tang Y, Dixon RA. 2011. MATE2 mediates vacuolar sequestration of flavonoid glycosides and glycoside malonates in Medicago truncatula. Plant Cell 23, 1536–1555.

Zhao J. 2015. Flavonoid transport mechanisms: how to go, and with whom. Trends Plant Science 20, 576–585.

